# Early life chronic stress-disrupted activity of the dorsal raphe nucleus selectively drives behavioral impairments

**DOI:** 10.1101/2025.08.05.668669

**Authors:** Zoltan K. Varga, Archana Golla, Florence Kermen

**Affiliations:** Department of Neuroscience, Faculty of Health and Medical sciences, University of Copenhagen, 2200, Copenhagen, Denmark; Department of Biology, Faculty of Natural Sciences, Norwegian University of Science and Technology, 7491, Trondheim, Norway

**Keywords:** chronic stress, early life stress, acute stress, dorsal raphe nucleus, serotonin, GABA, 2-photon calcium imaging, startle habituation, anxiety, zebrafish

## Abstract

Stress elicits variable systemic and neural changes in vertebrates, with outcomes ranging from adaptive to pathological. Several studies have implicated the dorsal raphe nucleus (DRN), a brainstem nucleus containing a heterogeneous population of serotonergic (5-HT) neurons, in the adaptive stress response and the pathological changes resulting from chronic stress. However, it is not known whether early life chronic stress affects the developing DRN activity, or whether the stress-induced changes affect 5-HT DRN neurons in a subregion- or phenotype-specific manner. To answer these questions, we used *in vivo* 2-photon calcium imaging of 5-HT DRN neurons in larval zebrafish exposed to chronic unpredictable stress during early life. We found that early life chronic stress prevented the normal habituation of the serotonergic system to a repeated acute stressor by altering the balance of excitatory/inhibitory responses within the DRN. Interestingly, these changes were most pronounced in a subset of stress-vulnerable serotonergic cells co-expressing GABAergic markers. Further, using chemogenetic ablation of 5-HT DRN neurons, we showed that stress-induced plasticity of the DRN contributed to changes in startle response habituation and in locomotive activity, but not in anxiety-like behaviors. Collectively, our results emphasize the role of stress-induced plasticity of DRN neurons in the selective regulation of maladaptive behavioral outcomes.

## Introduction

In vertebrates, postnatal neuronal circuits are plastic and undergo critical maturation processes, which are in part shaped by external factors ^1–3^. Early life stress, therefore, is an important risk factor that can result in long-term alterations in the structure and function of neural circuits underlying emotions, defensive behaviors, and the stress response ^4–9^. Epidemiology studies have established a strong link between postnatal stress exposure and anxiety disorders during adulthood in humans ^10–12^, warranting an increased understanding of how early-life stress affects the function of developing neural circuits.

Serotonergic neurons in the dorsal raphe nucleus (DRN) modulate the function of brain regions involved in emotions, defensive behaviors and the stress response ^13–17^ and are thought to play a key role in the development of stress-induced disorders during adulthood ^18–20^. Serotonin (5-HT) signaling and the activity of 5-HT DRN neurons are altered in acutely or chronically stressed animals ^4,9,21–24^. In turn, drugs affecting the levels of serotonin in the brain can alleviate the symptoms associated with mood disorders such as depression and anxiety ^25–28^. In addition, optogenetic and chemogenetic manipulations of serotonergic DRN neurons’ activity influence active coping, fear, and anxiety-like behaviors ^22,29–32^. Altogether, these findings point towards an important role for the serotonergic DRN in the adaptive and pathological stress response and suggest that it could also contribute to the long-term negative behavioral outcomes of early life stress. However, little is known regarding how early life stress affects the developing DRN during the sensitive postnatal period and how it impacts 5-HT DRN responses to subsequent adverse events, which could have long-term repercussions on adaptive stress responses later in life. Moreover, recent studies make it increasingly clear that serotonergic neurons are a heterogeneous neural population in terms of their connectivity ^17,33–36^, neurotransmitter phenotype ^34,37–39^ and function ^40,41^. In that context, whether stress differentially affects subpopulations of serotonergic DRN neurons in a region- or phenotype-specific manner is yet to be determined.

To answer these questions, we took advantage of the small size and transparency of larval zebrafish (*Danio rerio*), which allows for exploring the functional responses of the conserved 5-HT DRN neurons *in vivo* ^14,21,22,29,42^. We used 2-photon imaging in transgenic larval zebrafish expressing a calcium indicator in serotonergic neurons to investigate the effects of early life chronic unpredictable stress (CUS) on the number, phenotype, and functional response profile of 5-HT DRN neurons. Furthermore, we used chemogenetic ablation of 5-HT DRN neurons to determine their role in the behavioral consequences of chronic stress, with a particular focus on locomotion, startle habituation and anxiety.

## Methods

### Animals

The calcium imaging and the behavioral experiments were carried out in 154 transgenic zebrafish aged up to sixteen days. The following transgenic lines were used: Tg(*tph2:Gal4; uas:GCaMP6s;gad1b:DsRed*) ^14,43^ expressing the fluorescent calcium indicator GCaMP6s under the tryptophan hydroxylase 2 (Tph2) promoter in the dorsal raphe serotonergic neurons and the red fluorescent protein DsRed under the Glutamate decarboxylase 1 (Gad1b) promoter; Tg(*tph2:Gal4; uas:nitroreductase-mCherry*) expressing the bacterial nitroreductase (NTR) in serotonergic DRN neurons. Fish were raised at a constant temperature of 28°C, under a 12/12-hour (Norway) or 10/14-hour (Denmark) light cycle and fed twice a day with commercial flakes (TetraMin). Fish were housed in groups of 40-60 in meshed bottom nursery tanks in a recirculating system (Techniplast). The experiments were approved by the Norwegian Food Safety Authority (permit number: 17127) and the Danish Inspectorate of Animal experiment (permit number: 2023-15-0201-01493).

### Experimental design

#### Calcium imaging experiment

Fish were screened for GCaMP6s expression in the dorsal raphe nucleus and pan-neuronal DsRed expression at four-day post-fertilization (dpf) under an epifluorescence microscope (Axiolmager, ZEISS). At six dpf, fish were randomly assigned to control or stressed groups (protocol 1 and 2). Fish in the stressed group (protocol 1) were exposed to chronic unpredictable stress (CUS) from six to thirteen dpf (Fig 1A). An additional stressed group (protocol 2) was introduced that were not exposed to hyperosmotic shock as part of the CUS. *In vivo* calcium imaging was performed at 14 or 15 dpf.

**Figure 1:**
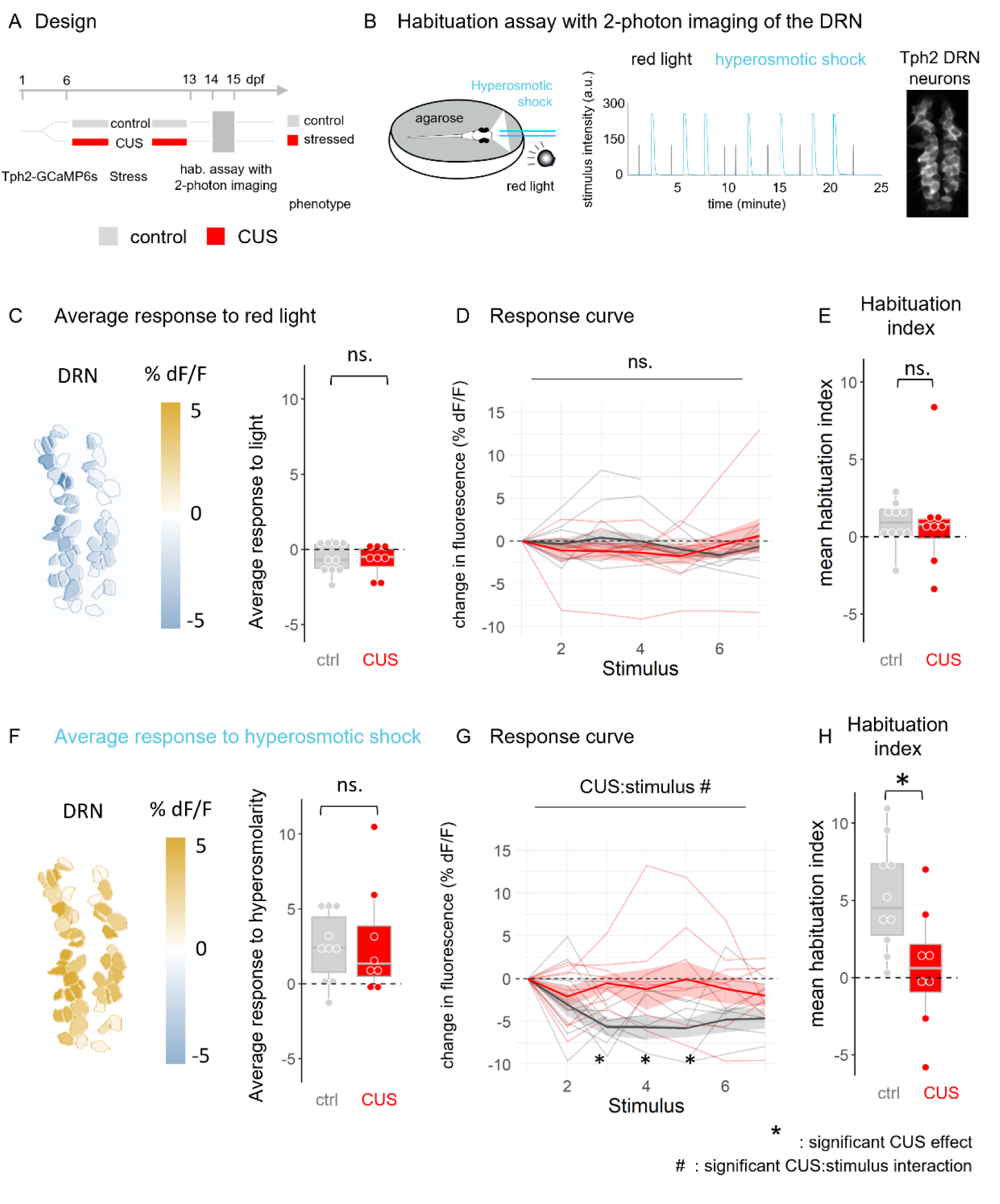
Light and hyperosmolarity responses of serotonergic neurons in the dorsal raphe nucleus. **A**. Schematic representation of the experimental design. **B**. Left. Scheme representing the experimental chamber holding 14-15 dpf Tg(*tph2:Gal4; uas:GCaMP6s; Gad1b:DsRed*) zebrafish larvae during 2-photon calcium imaging. Middle. Dynamics of light flashes (black) and hyperosmotic solution (cyan) stimuli during a typical recording. Right. Representative image of the DRN as recorded under the 2-photon microscope of a control fish (GCaMP6s labelling). **C**. Map of the average response of individual Tph2+ neurons to all red light flashes in the same fish as in B (left) and across all DRN neurons for each fish in the control and stressed groups (right, unpaired two-sided t-test: n (ctrl, cus) = 12,9; p = 0.488). **D**. Average change in the response amplitude of the DRN to repeated light stimuli relative to the first light presentation. Negative values indicate a decreased response; positive values indicate an increased response, 0 indicates no change (Repeated measure ANOVA: cus p = 0.281; stimulus number p = 0.808; cus:stimulus interaction p = 0.212). **E**. Mean of light response habituation indices (unpaired two-sided t-test: n (ctrl, cus) = 12,9; p = 0.900). **F**. Map of the average response of individual Tph2+ neurons to all hyperosmotic shocks in the same fish as in B (left) and across all DRN neurons for each fish in the control and stressed groups (right, unpaired two-sided t-test: n (ctrl, cus) = 12,9; p = 0.820). **G**. Average change in the response amplitude of the DRN to repeated hyperosmotic shocks relative to the first presentation (Repeated measure ANOVA: cus, p = 0.063; stimulus number, p = 0.007; cus:stimulus interaction, p = 0.030). **H**. Osmotic shock response habituation index (unpaired two-sided t-test: n (ctrl, cus) = 12,9; p = 0.022). Statistical parameters are shown in Table 1.

**Table 1:** CUS on DRN activation.

| Result / model | Test statistics<br>(t/F-values) | p-value | n |
| --- | --- | --- | --- |
| **average response to hyperosmolality** | *t* | *p* | Ctrl / CUS: 12 / 9 |
| *t.test(CUS)* | -0.233 | 0.820 |  |
| **mean habituation index** | *t* | *p* |  |
| *t.test(CUS)* | 2.572 | 0.022 * |  |
| **stimulus response to hyperosmolality** | *F* | *p* |  |
| *rmANOVA(CUS\*stimulus)* |  |  |  |
| CUS | 3.908 | 0.063 . |  |
| stimulus | 3.121 | 0.007 * |  |
| CUS*stimulus | 2.440 | 0.030 * |  |
| *post-hoc (control vs CUS)* | *t* | *p* |  |
| 2nd stim | -0.500 | 0.619 |  |
| 3rd stim | -2.723 | 0.009 * |  |
| 4th stim | -2.198 | 0.032 * |  |
| 5th stim | -2.761 | 0.008 * |  |
| 6th stim | -1.350 | 0.183 |  |
| 7th stim | -0.885 | 0.380 |  |
| **average response to light** | *t* | *p* |  |
| *t.test(CUS)* | 0.715 | 0.488 |  |
| **habituation index** | *t* | *p* |  |
| *t.test(CUS)* | -0.129 | 0.900 |  |
| **stimulus response to light** | *F* | *p* |  |
| *rmANOVA(CUS\*stimulus)* |  |  |  |
| CUS | 1.198 | 0.281 |  |
| stimulus | 0.059 | 0.808 |  |
| CUS*stimulus | 1.574 | 0.212 |  |

#### Behavioral phenotyping & DRN ablation experiments

The role of the DRN in CUS-induced behavioral changes was assessed using a battery of behavioral tests applied to four experimental groups (DRN intact control or CUS groups; DRN ablated control or CUS groups). Exposure to metronidazole (MTZ) for ablation and sham ablation took place after the end of the CUS protocol, at 14 dpf. Behavioral tests were conducted between 15 and 16 dpf. The habituation and the swimming plus-maze (SPM) assays were conducted in the same cohort of animals (n=113).

### Chronic unpredictable stress exposure

Fish in the stressed groups were exposed to two stressors per day that were applied at random times to maintain unpredictability, following a protocol previously established in our lab ^44^. The five following stressors were used: chasing (using a small net or a pipette for five minutes); turbulences (tank water replacements followed by increased air bubbling for three minutes, repeated three times); hyperosmotic shock (100 mM NaCl for ten minutes); light flashes exposure (6 mW/cm^2^ light flashes at 5Hz for ten minutes); pH drop (pH=4 for three minutes). Fish in the stressed group were exposed to the five above-mentioned stressors (protocol 1), while fish in the stressed group without prior hyperosmotic shock experience (protocol 2) were exposed to all stressors but hyperosmotic shock. Control groups were raised in the same conditions without stressor exposure but were briefly exposed to tank change every day, so they were accustomed to handling.

### *In vivo* 2-photon calcium imaging

Calcium imaging was performed in fish aged 14-15 dpf under a two-photon microscope (Leica SP8 inverted SMD/MP) equipped with a 25x water objective (Leica HCX IRAPO L). A mode-locked laser (Coherent Chameleon Vision-S) tuned to 1020 nm was used for fluorophore excitation, and band-pass emission filters centered around 525 nm and 640 nm were used for collecting images in the green and red channels, respectively. Volumetric imaging was done at 3.2Hz and 512 x 256-pixel resolution across three planes starting from the dorsal-most plane with visible Tph2+ neurons to 40µm ventrally, with 20µm distance between two planes. The fish were embedded ventral side up in 2% low-gelling temperature agarose (Sigma-Aldrich, CAT: A9414) in a glass-bottom petri dish (∼0.18 µm thick, VWR, CAT: 734-2904). Agarose was let to solidify for two minutes and the section covering the nose and mouth was carefully removed using a scalpel. The petri dish was then placed in a temperature-controlled chamber maintained at 28°C with constant air bubbling. For imaging, the embedded fish was placed on the microscope imaging platform and was supplied continuously with air-bubbled water delivered at the rate of 9 mL/minute and was let to further habituate for at least five minutes before the experiment started. Fish health was monitored before and after each recording by visual examination of the blood flow in the brain.

A 25-minute recording was obtained during which the fish was exposed to two different stimuli (red-light flashes and hyperosmotic solution). All fish were exposed to the same temporal sequence of stimuli, where stressors were applied seven times each, in pseudo-random order, with intervals of one minute after the red-light flashes and two minutes after the hyperosmotic solution. The light flashes were delivered at 5 Hz for 5 seconds using a red LED equipped with a long-pass filter (cut-off wavelength: 635 nm, Thorlabs) positioned in front of the fish. The hyperosmotic solution (100 mM NaCl) was bubbled with air and applied via the perfusion tube to the fish’s nose using a solenoid switch valve (three-way valve, Bio-Chem Fluidics), which opened for five seconds. Stimuli delivery was automatically triggered using a programmed microcontroller board (Arduino Uno). To verify the timing of light flashes and hyperosmotic solution delivery in the recording chamber, an additional 25-minute recording was performed at the end of the recording session after replacing the hyperosmolarity solution with a 1.5 mM fluorescein solution. While light flashes lasted five seconds, the gradual diffusion of hyperosmolarity stimulus in water caused the fluorescein concentration to peak in front of the fish approximately 15 seconds after the onset.

### Calcium imaging analysis

Consecutive frames were aligned using the FIJI TurboReg plugin ^45^ to correct for tissue drift and fish movement. Aligned frames were visually inspected, and recordings with persisting tissue drift were discarded. Only recordings with three or more repetitions of both stimuli were further analyzed. Tph2+ neurons were manually segmented in Matlab, and the resulting neuron mask was overlaid on the corresponding red channel average for double-positive (Tph2+ Gad1b+) neurons detection. Pixel intensity in each neuron was averaged for each time frame, and the relative change in fluorescence (% dF/F_0_) was calculated as follows:

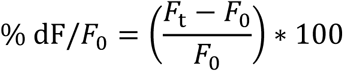

in which F_0_ was the average fluorescence for each neuron during the pre-stimulus baseline of 15 seconds, and F_t_ was the fluorescence of the same neuron at time frame t. The response to stimuli was then calculated by averaging %dF/F_0_ during the period following the stimulus onset (five seconds for light flashes and 15 seconds for hyperosmotic solution). Neurons were categorized into excited, inhibited or non-responsive in Figure 2, when the stimulus amplitude response was > +1%dF/F_0_, < - 1%dF/F_0_, or between −1% and 1%dF/F_0,_ respectively. The relative proportions of inhibitory and excitatory responses to successive hyperosmotic shock presentations (Figure 2F-H) was calculated as follows: (E-I)/(E+I), where E is the proportion of excited neurons and I is the proportion of inhibited neurons. Mean habituation index was averaged from habituation indices (H) of stimulus responses. Hs were calculated by subtracting the hyperosmotic response of each stimulus (2^nd^, 3^rd^, 4^th^, 5^th^, 6^th^, 7^th^) from the response value of the first stimulus.

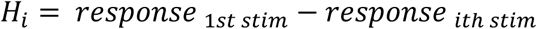

, where response 1^st^ stim is the response to the first stimulus and response_ith_ is the response to the i^th^ stimulus. Habituation of the response is therefore indicated by positive habituation indices.

**Figure 2.**
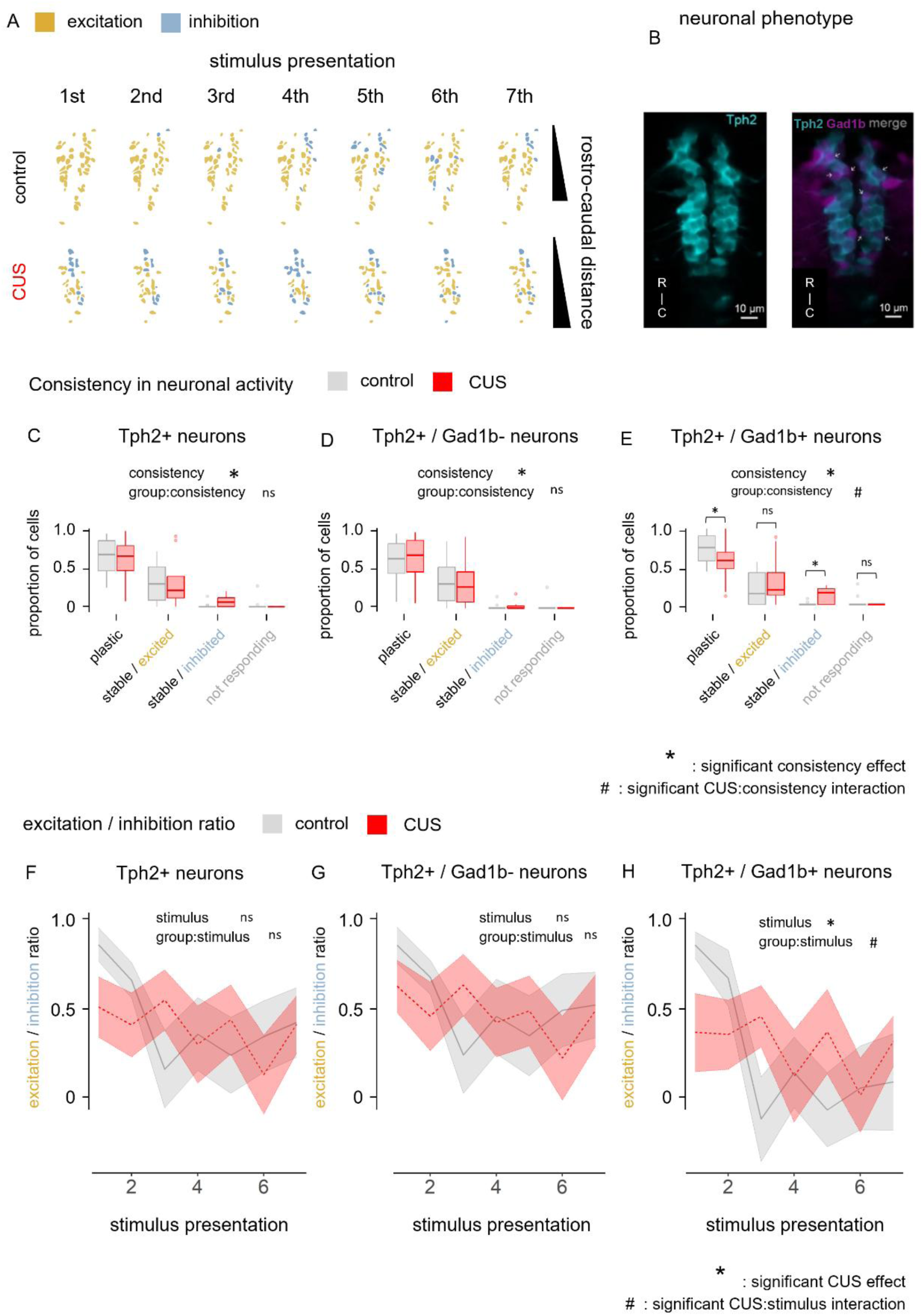
Chronic early life stress alters the balance of excitatory/inhibitory responses in the DRN in a cell-type specific manner. **A.** Maps of DRN responses to all hyperosmotic shock presentations (left to right: 1^st^ to 7^th^) in two representative control and CUS-treated fish (gold: excitation, blue: inhibition). **B.** Two-photon image of the dorsal raphe nucleus in a 14 dpf *Tg(tph2:Gal4; uas:GCaMP6s; Gad1b:DsRed)* zebrafish illustrating that a subpopulation of Tph2 (cyan) neurons, indicated by white arrows, co-expresses Gad1b (magenta). **C-E.** Consistency of neuronal responses in all serotonergic DRN neurons (C)(ANOVA: consistency p = 1.275e-15, cus:consistency p = 0.782), Gad1b-serotonergic neurons (D)(ANOVA: consistency p = 7.145e-14, cus:consistency p = 0.992) and Gad1b+ serotonergic neurons (E)(ANOVA: consistency p < 2e-16, cus:consistency p = 0.021; pairwise posthoc (t.test/brunner-munzel with FDR adjustment): plastic p = 0.016, stable excited p = 0.121, stable inhibited p = 0.049, non-responding p = 0.694). Stable neurons remained excited, or inhibited, throughout the experiment (in response to 6 out of 7 presentations). Plastic neurons switched between excited and inhibited responses. **F-H.** Relative proportions of inhibitory and excitatory responses (see methods) to successive hyperosmotic shock presentations in all serotonergic DRN neurons (F)(repeated measure ANOVA: cus p = 0.870, stimulus number p = 0.015, cus:stimulus number p = 0.070), Gad1b-serotonergic neurons (G)(repeated measure ANOVA: cus p = 0.918 stimulus number p = 0.036, cus:stimulus number p = 0.119) and Gad1b+ serotonergic neurons (H)(repeated measure ANOVA: cus p = 0.473, stimulus number p = 0.005, cus:stimulus number p = 0.036). Values close to 1 indicate mostly excitatory responses in the DRN; values around 0 indicate similar proportions of excitatory and inhibitory responses; negative values indicate more inhibitory than excitatory responses. n (ctrl, cus) = 12, 9. Statistical parameters are shown in Table 2 and Supplementary Table 1.

**Table 2:**
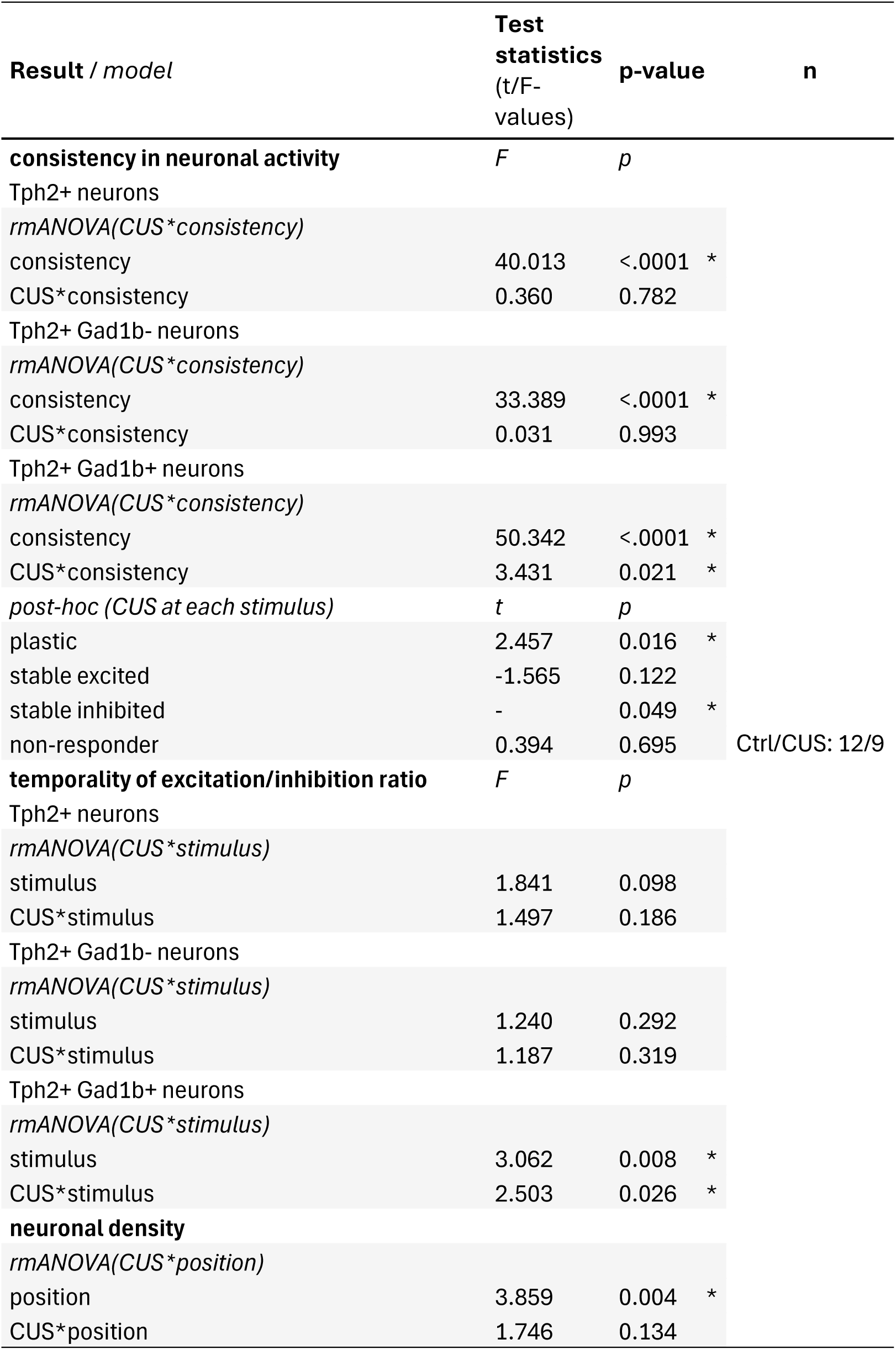
CUS on DRN activation (temporal, spatial and phenotypic dimensions)

### Serotonergic DRN neuron ablation

For ablation of serotonergic DRN neurons, metronidazole (5 mM MTZ in AFW; Merck) was applied to nitroreductase-expressing *Tg(tph2:Gal4:UAS-ntr-mCherry)* fish, and to their non-expressing siblings (mCherry negative; sham ablation). All fish were kept in aluminum foil-covered petri dishes during treatment to prevent degradation of light-sensitive metronidazole. The 24-hour treatment started in the morning of 14 dpf and was followed by a 24-hour recovery period before behavioral testing. The ablation of serotonergic DRN neurons was verified by euthanizing larvae after the behavioral assay, fixing them overnight in formalin 10% with 0.25% TritonX, and scanning the brains under a confocal microscope (LSM 700 AxioImager 2, Zeiss) equipped with 10× and 40× water-immersion objectives (Supp. Fig 3). For visualization of Tph2+ neurons, four additional *Tg(tph2:Gal4:UAS-ntr-mCherry)* fish positive for mCherry were exposed to vehicle and euthanized at 16 dpf.

### Behavioral assays & analysis

All behavioral recordings were performed during the second half of the light period in a temperature-controlled cabinet (28°C). Fish trajectory was detected off-line on the recorded videos using Ethovision (Noldus).

#### Habituation assay

The assays were conducted in standard 24-well plates in a custom-made setup. The recording took place in a dark room, where the plate was placed on a horizontal support enabling homogeneous illumination from below, using a white LED strip and an opaque Plexiglas sheet as a diffuser. Additionally, an infrared LED strip was placed under the plate to allow for recording and detection of animal movement during the dark flashes, regardless of the lack of visible light. Recording of the behaviour was done at 5 frames per second using a camera (Basler ace2) placed 30 cm above the arena and equipped with a lens (FUJIAN; focal length=35 mm) covered with a long-pass infrared filter. Dark flashes were delivered automatically using an Arduino coupled to a relay that switched the white LED strip on and off.

Fish from the four experimental groups were distributed in each well of the plates in randomized order. A 10-minute baseline recording period with constant illumination was followed by a 5-minute stimulus period consisting of 1-second-long dark flashes repeated 50 times at 5-second inter-stimulus-intervals (ISI). Such stimulus presentation was selected based on previously published habituation protocols (Pantoja 2016). *Habituation assay analysis*. Instantaneous distances swam were extracted using Ethovision and responses to all dark flashes were calculated as follows:

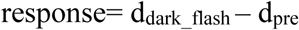

where d_dark_flash_ is the average displacement during the 1 sec of dark flash and d_pre_ is the average displacement during the 1 sec prior to the dark flash. For display in some panels of Figure 3 and Supplementary Figure 3, the responses to 3 consecutive dark flashes were averaged into stimulus bins. Habituation indices (H) were calculated by subtracting the dark flash response of a particular stimulus bin (2nd, 4th, 6th, 8th, 10th, or 16th) from the response value of the first stimulus bin.

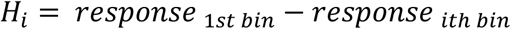

, where response_1st bin_ is the average response to the first stimulus bin of 3 dark flashes and response_ith bin_ is the average response to the i^th^ stimulus bin of 3 dark flashes. Habituation of the response is therefore indicated by positive habituation indices. Correlation between first response bins and habituation indices were calculated within each group using Pearson’s correlations.

**Figure 3:**
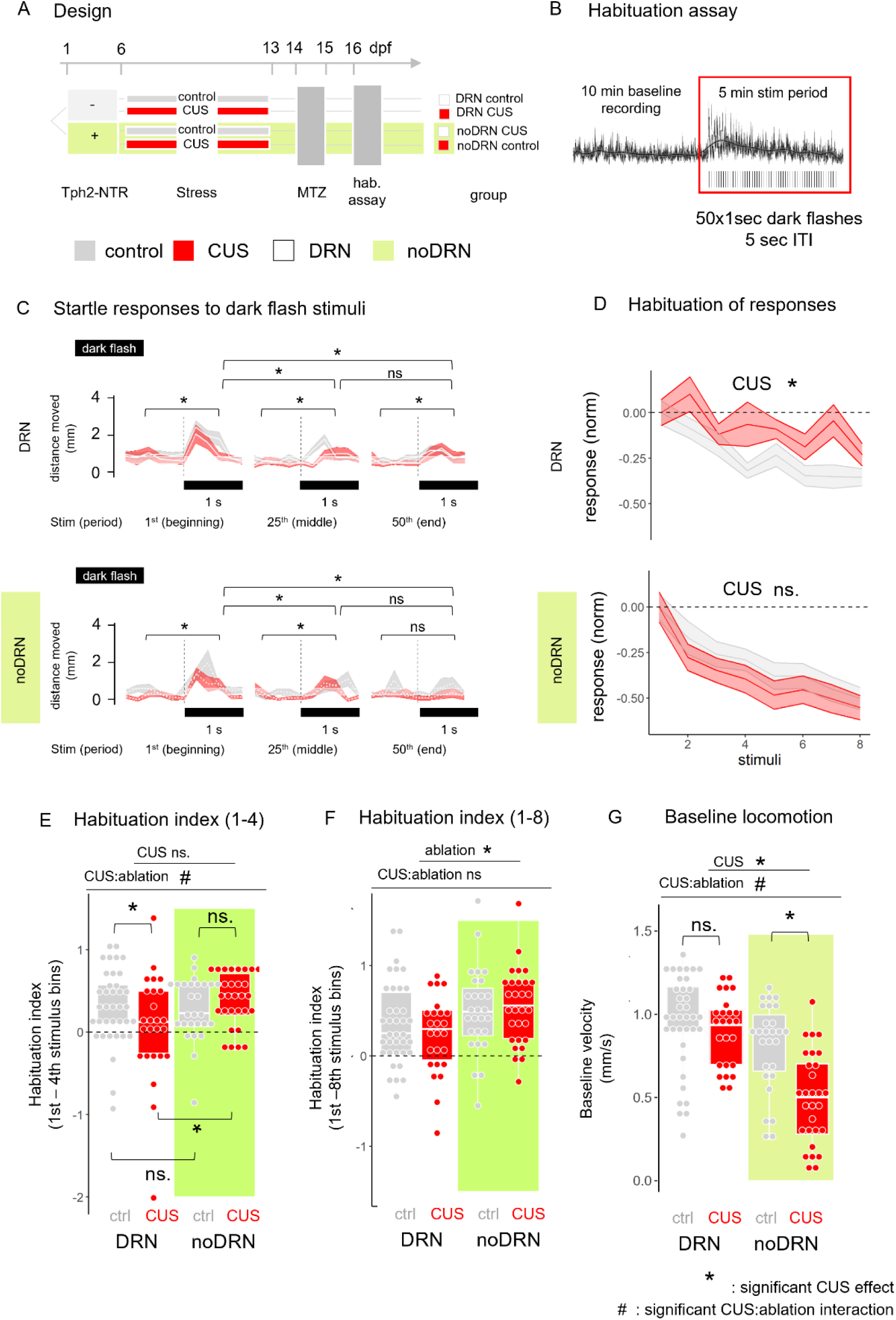
Chronic early life stress disrupts behavioral stress-habituation through serotonergic DRN function. **A.** Schematic representation of the experimental design. Larvae were sorted to nitroreductase (NTR) expressing (+) and non-expressing (−) subjects, then allocated to control or CUS treatment. Metronidazole (MTZ) treatment and a habituation assay were done to all subjects. **B.** A representative image to show the stages and protocol of the habituation assay: 10 minutes of baseline recording were followed by a 5-minute stimulus period that consisted of 50 1-second-long dark flashes separated by 5-second inter-stimulus-intervals (ISIs). **C.** Startle responses to dark flashes at the beginning (1^st^ dark flash), middle (25^th^ dark flash), and the end (50^th^ dark flash) of the stimulus period in each group. Dark flash timing and duration is indicated by black bar. Statistical comparison of pre-stimulus and stimulus periods within and across dark flash presentations is indicated (* p < 0.05). **D.** Habituation of responses to dark flash presentations across groups. The response to dark flashes (average distance moved during dark flash - average distance moved during 1 s before) was averaged for 3 consecutive dark flashes (stimulus bins) and normalized to the first stimulus bin. The first 8 stimulus bins are presented. **E.** Response habituation index between the 1^st^ and the 4^th^ stimulus bins. **F.** Response habituation index between the 1^st^ and the 8^th^ stimulus bins. **G.** Mean swimming velocity during the baseline recording period (no stimuli). Asterisk indicates CUS main effect, hashtag indicates CUS:ablation interactions. Statistical results for all panels are available in Table 3. n (DRN ctrl, DRN cus, noDRN ctrl, noDRN cus) = 38, 27, 27, 29.

**Table 3:**
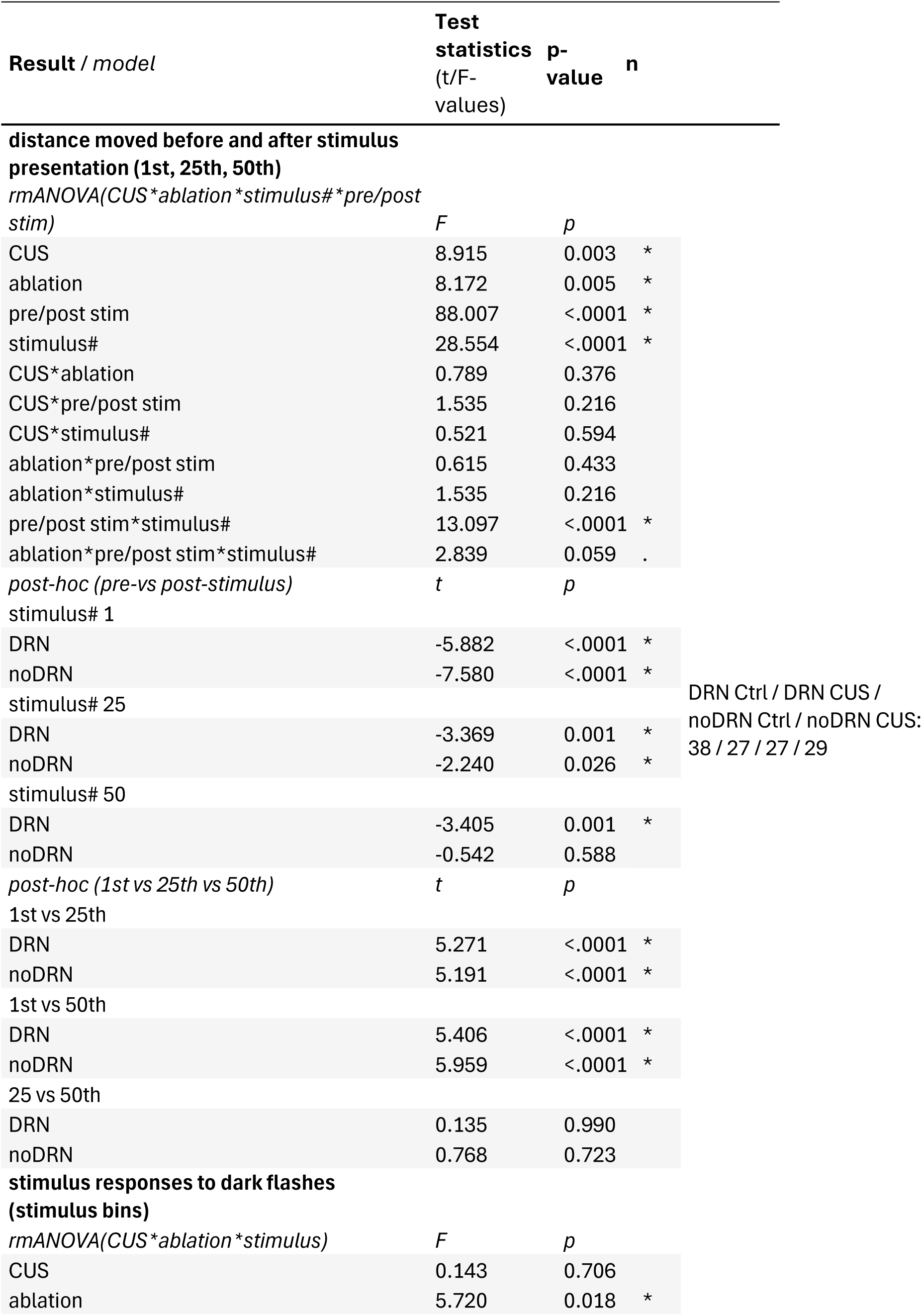
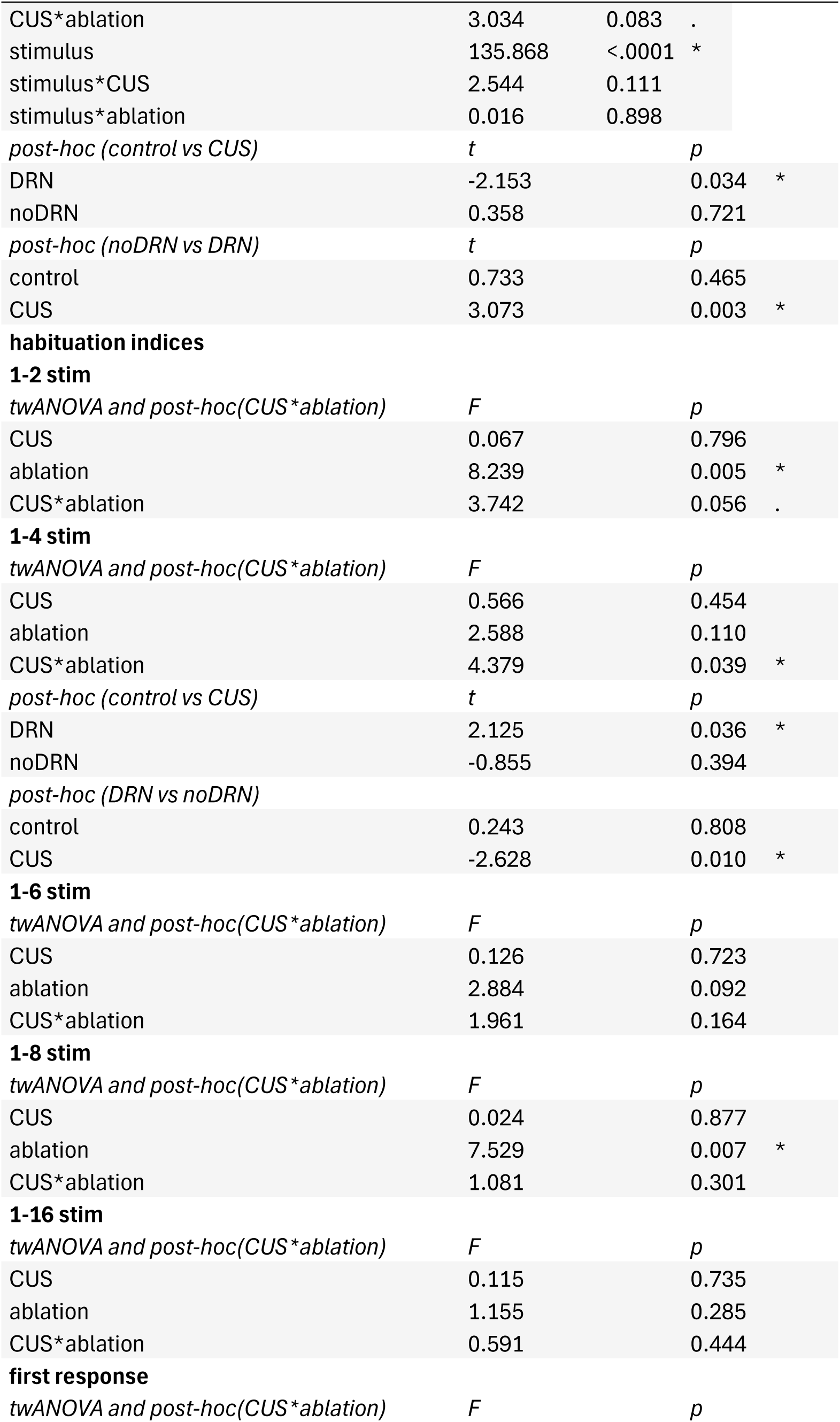
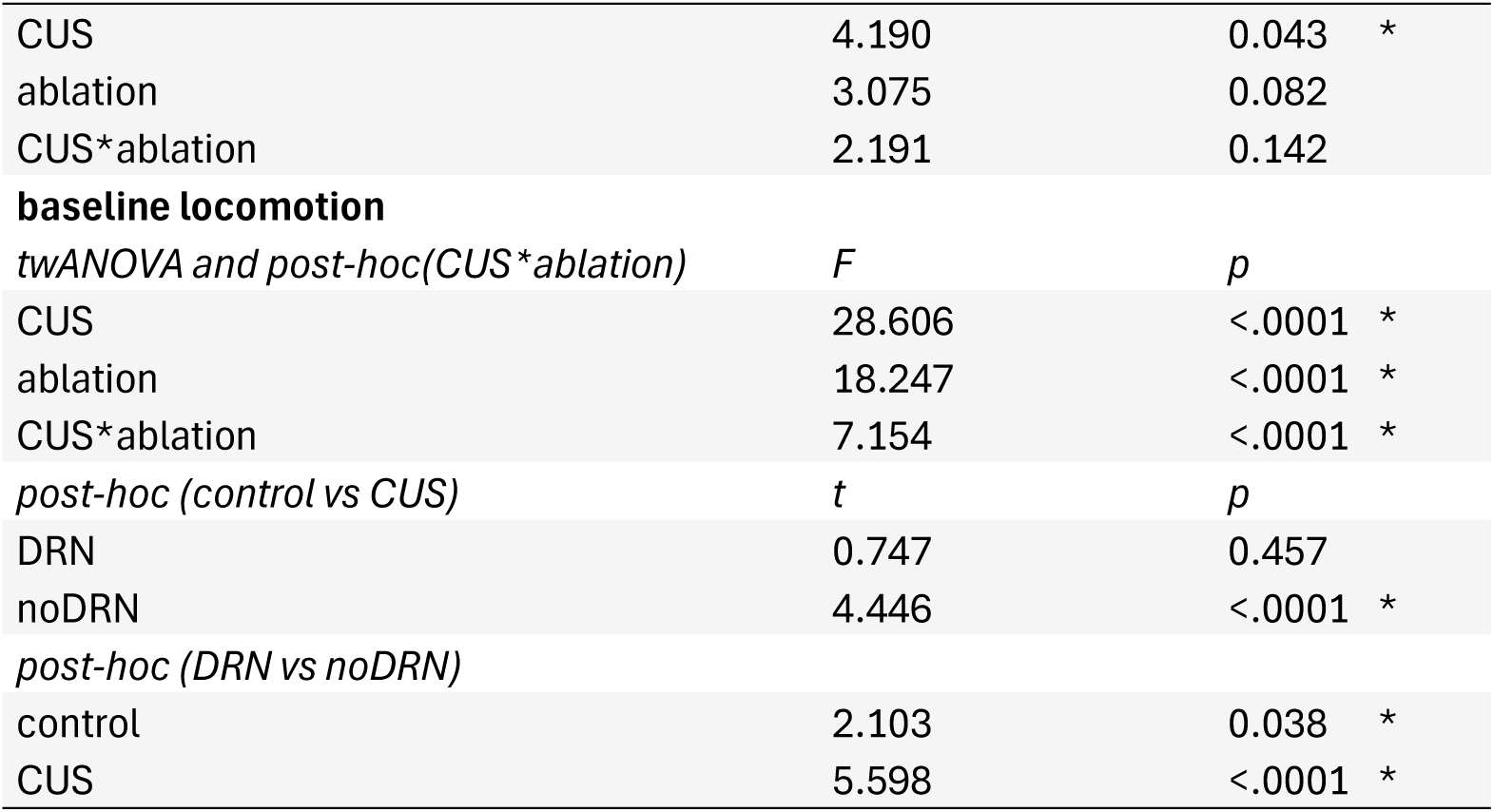
CUS and DRN ablation on behavioural habituation and locomotion.

| **Result / model** | **Test statistics (t/F-values)** | **p-value** | **n** |  |
| --- | --- | --- | --- | --- |
| **distance moved before and after stimulus presentation (1st, 25th, 50th)** |  |  |  |  |
| *rmANOVA(CUS\*ablation\*stimulus#\*pre/post stim)* | *F* | *p* |  |  |
| CUS | 8.915 | 0.003 | * |  |
| ablation | 8.172 | 0.005 | * |  |
| pre/post stim | 88.007 | <.0001 | * |  |
| stimulus# | 28.554 | <.0001 | * |  |
| CUS*ablation | 0.789 | 0.376 |  |  |
| CUS*pre/post stim | 1.535 | 0.216 |  |  |
| CUS*stimulus# | 0.521 | 0.594 |  |  |
| ablation*pre/post stim | 0.615 | 0.433 |  |  |
| ablation*stimulus# | 1.535 | 0.216 |  |  |
| pre/post stim*stimulus# | 13.097 | <.0001 | * |  |
| ablation*pre/post stim*stimulus# | 2.839 | 0.059 | . |  |
| *post-hoc (pre-vs post-stimulus)* | *t* | *p* |  |  |
| stimulus# 1 |  |  |  |  |
| DRN | -5.882 | <.0001 | * |  |
| noDRN | -7.580 | <.0001 | * |  |
| stimulus# 25 |  |  |  | DRN Ctrl / DRN CUS /<br>noDRN Ctrl / noDRN CUS:<br>38 / 27 / 27 / 29 |
| DRN | -3.369 | 0.001 | * |  |
| noDRN | -2.240 | 0.026 | * |  |
| stimulus# 50 |  |  |  |  |
| DRN | -3.405 | 0.001 | * |  |
| noDRN | -0.542 | 0.588 |  |  |
| *post-hoc (1st vs 25th vs 50th)* | *t* | *p* |  |  |
| 1st vs 25th |  |  |  |  |
| DRN | 5.271 | <.0001 | * |  |
| noDRN | 5.191 | <.0001 | * |  |
| 1st vs 50th |  |  |  |  |
| DRN | 5.406 | <.0001 | * |  |
| noDRN | 5.959 | <.0001 | * |  |
| 25 vs 50th |  |  |  |  |
| DRN | 0.135 | 0.990 |  |  |
| noDRN | 0.768 | 0.723 |  |  |
| **stimulus responses to dark flashes (stimulus bins)** |  |  |  |  |
| *rmANOVA(CUS\*ablation\*stimulus)* | *F* | *p* |  |  |
| CUS | 0.143 | 0.706 |  |  |
| ablation | 5.720 | 0.018 | * |  |

|  |  |  |  |
| --- | --- | --- | --- |
| CUS*ablation | 3.034 | 0.083 | . |
| stimulus | 135.868 | <.0001 | * |
| stimulus*CUS | 2.544 | 0.111 |  |
| stimulus*ablation | 0.016 | 0.898 |  |
| *post-hoc (control vs CUS)* | *t* | *p* |  |
| DRN | -2.153 | 0.034 | * |
| noDRN | 0.358 | 0.721 |  |
| *post-hoc (noDRN vs DRN)* | *t* | *p* |  |
| control | 0.733 | 0.465 |  |
| CUS | 3.073 | 0.003 | * |
| **habituation indices** |  |  |  |
| **1-2 stim** |  |  |  |
| *twANOVA and post-hoc(CUS\*ablation)* | *F* | *p* |  |
| CUS | 0.067 | 0.796 |  |
| ablation | 8.239 | 0.005 | * |
| CUS*ablation | 3.742 | 0.056 | . |
| **1-4 stim** |  |  |  |
| *twANOVA and post-hoc(CUS\*ablation)* | *F* | *p* |  |
| CUS | 0.566 | 0.454 |  |
| ablation | 2.588 | 0.110 |  |
| CUS*ablation | 4.379 | 0.039 | * |
| *post-hoc (control vs CUS)* | *t* | *p* |  |
| DRN | 2.125 | 0.036 | * |
| noDRN | -0.855 | 0.394 |  |
| *post-hoc (DRN vs noDRN)* |  |  |  |
| control | 0.243 | 0.808 |  |
| CUS | -2.628 | 0.010 | * |
| **1-6 stim** |  |  |  |
| *twANOVA and post-hoc(CUS\*ablation)* | *F* | *p* |  |
| CUS | 0.126 | 0.723 |  |
| ablation | 2.884 | 0.092 |  |
| CUS*ablation | 1.961 | 0.164 |  |
| **1-8 stim** |  |  |  |
| *twANOVA and post-hoc(CUS\*ablation)* | *F* | *p* |  |
| CUS | 0.024 | 0.877 |  |
| ablation | 7.529 | 0.007 | * |
| CUS*ablation | 1.081 | 0.301 |  |
| **1-16 stim** |  |  |  |
| *twANOVA and post-hoc(CUS\*ablation)* | *F* | *p* |  |
| CUS | 0.115 | 0.735 |  |
| ablation | 1.155 | 0.285 |  |
| CUS*ablation | 0.591 | 0.444 |  |
| **first response** |  |  |  |
| *twANOVA and post-hoc(CUS\*ablation)* | *F* | *p* |  |

|  |  |  |
| --- | --- | --- |
| CUS | 4.190 | 0.043 * |
| ablation | 3.075 | 0.082 |
| CUS*ablation | 2.191 | 0.142 |
| **baseline locomotion** |  |  |
| *twANOVA and post-hoc(CUS\*ablation)* | *F* | *p* |
| CUS | 28.606 | <.0001 * |
| ablation | 18.247 | <.0001 * |
| CUS*ablation | 7.154 | <.0001 * |
| *post-hoc (control vs CUS)* | *t* | *p* |
| DRN | 0.747 | 0.457 |
| noDRN | 4.446 | <.0001 * |
| *post-hoc (DRN vs noDRN)* |  |  |
| control | 2.103 | 0.038 * |
| CUS | 5.598 | <.0001 * |

#### Swimming plus-maze test (SPM)

The swimming plus-maze test developed by Varga et al. ^46^ was used to measure anxiety-like responses. The test apparatus was custom made using a 3D printer and consisted of two shallow arms (L x W x D: 10 x 8 x 2.5 mm) and two deep arms (L x W x D: 10 x 8 x 5 mm) separated by a center zone (L x W x D: 8 x 8 x 5 mm). The same camera and illumination were used as for the habituation setup above. Fish were pipetted from their home-tank to a well-plate, transferred to the setup and placed in the center zone of the apparatus. Fish movement was recorded for 6 minutes. The time spent in the shallow arms and the mean swimming velocity were measured.

### Statistical analysis

Data was evaluated using R statistical environment (R core team). 1.5 inter-quartile range (IQR) discrimination method was used on the whole experimental populations to identify and exclude potential outliers. Data including two groups were compared using student’s t-tests or the Brunner Munzel test (generalized Wilcoxon test) in case of uneven variability between compared groups. Data including repeated factors such as data points from the same animals at different positions (distance from the rostral-most cell) or at different time points of a test (habituation responses at the cellular or behavioral level) were analyzed with repeated measure ANOVA (rmANOVA). Data including multiple factors were analyzed with two-way interactional ANOVA (twANOVA). In the case of significant interactions between factors, post-hoc pairwise comparisons (PHC) were performed. In PHCs following rmANOVA, contrasts were only made between the first and the remaining levels and p-values were adjusted using the Dunett method. In PHCs following twANOVA, contrasts were only made within separate factor levels, e.g. assessing the difference between the non-stressed and stressed groups at the levels of non-ablated and ablated DRN to avoid unnecessarily high number of comparisons and p-values were adjusted using the false-discovery-rate (FDR) method. Data were reported as mean ± standard error of the mean (SEM) in the text and figure legends. A p-value < 0.05 was considered significant. All statistical results are shown in Tables 1-4 and Supplementary Table 1.

## Results

### Chronic early life stress alters the habituation of serotonergic neurons to hyperosmolarity stress

Previous studies have shown that the activity of 5-HT DRN neurons is modulated during a broad range of sensory-motor responses ^14,29,42^, including those evoked by aversive or stressful stimuli ^21,22,29^. However, it is not known how 5-HT DRN responses to aversive stimuli are affected by prior developmental stress exposure. To determine the effect of chronic early life stress on the DRN, we exposed Tg(*tph2:Gal4; uas:GCaMP6s; Gad1b:DsRed*) zebrafish to mild unpredictable stressors from 6-13 (dpf) and compared the DRN responses to two types of repeated stimuli (flashes of red light and high salinity solution) to that in control unstressed groups (Fig 1A-B). Flashes of red light were of low intensity and likely provided a neutral sensory stimulation, whereas high salinity solution is a well-documented aversive stimulus (hyperosmotic shock) that activates the HPI axis in zebrafish ^47–49^.

Overall, we found that the DRN did not react to light flashes but was excited by hyperosmotic solution, when the responses to all stimulus presentations were averaged across neurons (Fig 1C, F). There was no difference in these average responses between control and stressed groups. The initial response of the DRN to hyperosmotic solution or light flashes was also similar between control and stressed groups (Supp Fig 1A-B).

To examine the DRN response’s dynamic to subsequent presentations, we then plotted the difference in amplitude between the first response and that to subsequent stimuli (Fig 1D, G). We found no habituation of the DRN responses to light flashes (Fig1D) and no difference in habituation index between the control and stressed groups (Fig 1E). Conversely, the amplitude of the hyperosmolarity responses decreased across stimulus presentations in control, but not in stressed fish (Fig 1G). Comparing the response habituation index between groups confirmed that DRN hyperosmolarity responses habituated significantly less in stressed than in control fish (Fig 1H).

One potential confounder in interpreting those results is that stressed fish have been exposed to hyperosmolarity solution during the CUS protocol, and control fish have not. Therefore, at the time of measuring the DRN responses, the two groups differ in terms of early life experience and in familiarity with the hyperosmolarity solution presented acutely. To determine whether familiarity might contribute to the difference in habituation between the groups, we also used a CUS protocol without hyperosmotic shock (protocol 2) and found DRN neurons’ responses to hyperosmotic stimulus did not habituate in those fish (Supp Fig 1C-D). This suggests that the impaired habituation of DRN neurons measured in both stressed groups is caused by a prior history of early life adversity, rather than by the familiarity with the stimulus. Statistical parameters are shown in Table 1.

### Chronic early life stress alters the balance of excitatory/inhibitory responses in the DRN in a cell-type specific manner

Next, we sought to investigate the changes in neural circuit activity that might underlie the blunted DRN habituation in the CUS-exposed animals. Visual inspection of activity maps revealed that the 5-HT DRN neurons displayed heterogeneous responses to hyperosmotic shock, with a combination of excited and inhibited neurons (Fig 2A). The amount of excitatory and inhibitory responses appeared to be dynamic across stimulus presentations in control fish, and less so in the stressed group. This raised the interesting possibility that habituation in the control fish’s DRN might be partly driven by a change in the relative proportions of excitatory and inhibitory responses.

To test this hypothesis, we first quantified the consistency of excitatory and inhibitory responses of individual neurons throughout stimulus presentations (Fig 2C-E). Most serotonergic neurons exhibited a mixed response profile and alternated excitatory and inhibitory responses (plastic responses); whereas other neurons remained consistently excited, or consistently inhibited, throughout the recording (stable responses). There was no interaction between chronic early life stress exposure and the consistency of responses when assessing all serotonergic neurons together (Fig 2C). However, chronic stress significantly affected the consistency of responses within a subset of serotonergic neurons that co-express a GABAergic marker (Gad1b, Fig 2E). Stressed fish exhibited a significantly reduced proportion of plastic Tph2 Gad1b+ neurons and an increased proportion of stable-inhibited Tph2 Gad1b+ neurons, compared to controls (Fig 2E). This was not due to a direct effect of CUS on the number of Tph2+ neurons (Sup. Fig 2D) or on the proportion of double-positive neurons, which remained unchanged in control and stressed groups (Sup. Fig 2C). Interestingly, while plastic neurons – regardless of their phenotype – show similar spatial distribution in both groups, stably inhibited Gad1b+ cells in CUS-exposed fish tend to have a more caudal location in the DRN (Sup. Fig 2E). This is particularly interesting, given that Gad1b+ cells are more abundant in the rostral part of the DRN in general (Sup. Fig. 2B).

Next, we examined the change in excitatory/inhibitory response balance throughout the recording and within the different subsets of serotonergic cells (Fig 2F-2H). Overall, in the control group, the balance shifted towards a decreased proportion of excitatory relative to inhibitory responses in the DRN during the course of habituation (Fig 2F). This shift was less pronounced in the stressed group, in line with the blunted habituation in those animals. Interestingly, the difference in temporal dynamics of the excitatory/inhibitory balance between stressed and control groups was again most apparent in the Gad1b+ subset of serotonergic cells (Fig 2H).

Altogether, we showed that chronic stress impaired the habituation of serotonergic neurons’ response to a repeated acute stressor. This impaired habituation in CUS-exposed fish was a result of the lack of plasticity of neurons – Gad1b+ serotonergic cells in particular – to switch from excitation to inhibition in response to the repeated acute stress. Statistical parameters are shown in Table 2 and Supplementary Table 1.

### Chronic stress disrupts behavioral stress-habituation through serotonergic DRN function

To determine whether CUS-induced impairment of DRN 5-HT neurons’ habituation translates into impairment of habituation at the behavioral level, we measured habituation of locomotion in control and CUS fish in response to a series of aversive dark flashes. In addition, we tested whether 5-HT DRN neurons played a causal role in driving changes in behavioral habituation by assessing the performance of DRN intact (Ntr-, exposed to MTZ) and DRN ablated (Ntr+, exposed to MTZ) fish (Fig 3A). MTZ exposure was performed 2 days before behavioral testing and successfully ablated most Tph2+ neurons in the DRN of Tph2:Gal4:UAS:Ntr-mCherry+ fish (Sup. fig 3 A-B). The movement of all fish was recorded for 10 minutes in the light (baseline period), followed by 5-minute exposure to 1-second dark flashes (50 flashes at 5-second intervals; Fig 3B, Sup. fig 4A).

Fish initially showed a clear startle response to the dark flashes in all groups, as they responded by increasing the distance swam (Fig 3C, left). The comparison of distance swam during the 1^st^, 25^th^ and last dark flash shows that the amplitude of the startle response is substantially decreased mid-experiment and remains low until the last dark flash in all groups (Fig 3C: left to right; Sup. Fig 4A), indicating that the habituation already happened at the first part of the test.

To handle natural variability in startle responses and facilitate visualization, we averaged responses to three consecutive dark flashes (binned responses) and focused the rest of the analysis on the first half of the stimulus period, i.e. the 8 binned responses corresponding to 24 dark flashes (Fig 3D). Distance swam in the whole stimulus period, pre-and post-stimulus periods as well as responses to each stimulus are shown in both 1 second resolution and 1 stimulus bin resolution in Supplementary figure 4A and 4B, respectively. Time dynamics of each response in the entire test are shown in supplementary figure 4. Regardless of the resolution of the analysis, we found that zebrafish with an intact DRN quickly decreased their locomotor response to dark flashes indicative of habituation, and that previous CUS exposure significantly impaired this habituation (Fig 3D top). In contrast, both control- and CUS-exposed DRN-ablated groups habituated to dark flashes, which indicates that DRN ablation rescues CUS-induced deficit in habituation (Fig 3D bottom). The habituation index, which quantifies the amplitude of the decrease in startle response relative to the beginning of the assay, revealed that the rescue effect of ablation is apparent in the first few stimulus bins (Fig 3D-3E, Sup. fig 4C). Besides these interactions, we found a general habituation-increasing effect of ablation at the end of the first part of the test (8^th^ stimulus bins, Fig 3F). During the second part of the test, we did not detect significant interaction between CUS and ablation anymore, however, the non-ablated CUS-exposed group showed close-to-zero habituation indices in the whole course of the test (Sup. Fig 4C).

CUS also had a main effect on the amplitude of the response to the first dark flash, with CUS fish displaying lower responses (Sup. Fig 4D). To verify that the lower initial reactivity of CUS-exposed fish is not what drives the decreased habituation in this group, we correlated the habituation indices with the magnitude of the first responses (Sup. Fig 4E). We found that the level of habituation correlated with the magnitude of the first response in all but the DRN intact CUS group, which confirmed that this group expresses different habituation dynamics which is independent from the magnitude of their first response (Sup. Fig 4E).

CUS exposure also had a general main effect and an interactional effect with ablation on locomotion during the baseline recording period (Figure 3G). Post-hoc comparisons revealed that CUS exposure selectively decreased locomotion in DRN-ablated animals.

In summary, we found that chronic early life stress impaired both neuronal- and behavioral-level habituation. Moreover, behavioral habituation could be rescued by the ablation of serotonergic neurons of the dorsal raphe nucleus. Statistical results for all panels are available in Table 3.

### Chronic stress enhances anxiety-like behavior independently from serotonergic DRN function

Given the well-established association between chronic stress and anxiety ^50^, we sought to investigate the role of the serotonergic DRN in anxiety-like behavior in CUS-exposed zebrafish. We applied the same experimental design as described above for the habituation but assessed anxiety-like responses in the swimming plus-maze (SPM, Fig 4 A, B).

**Figure 4:**
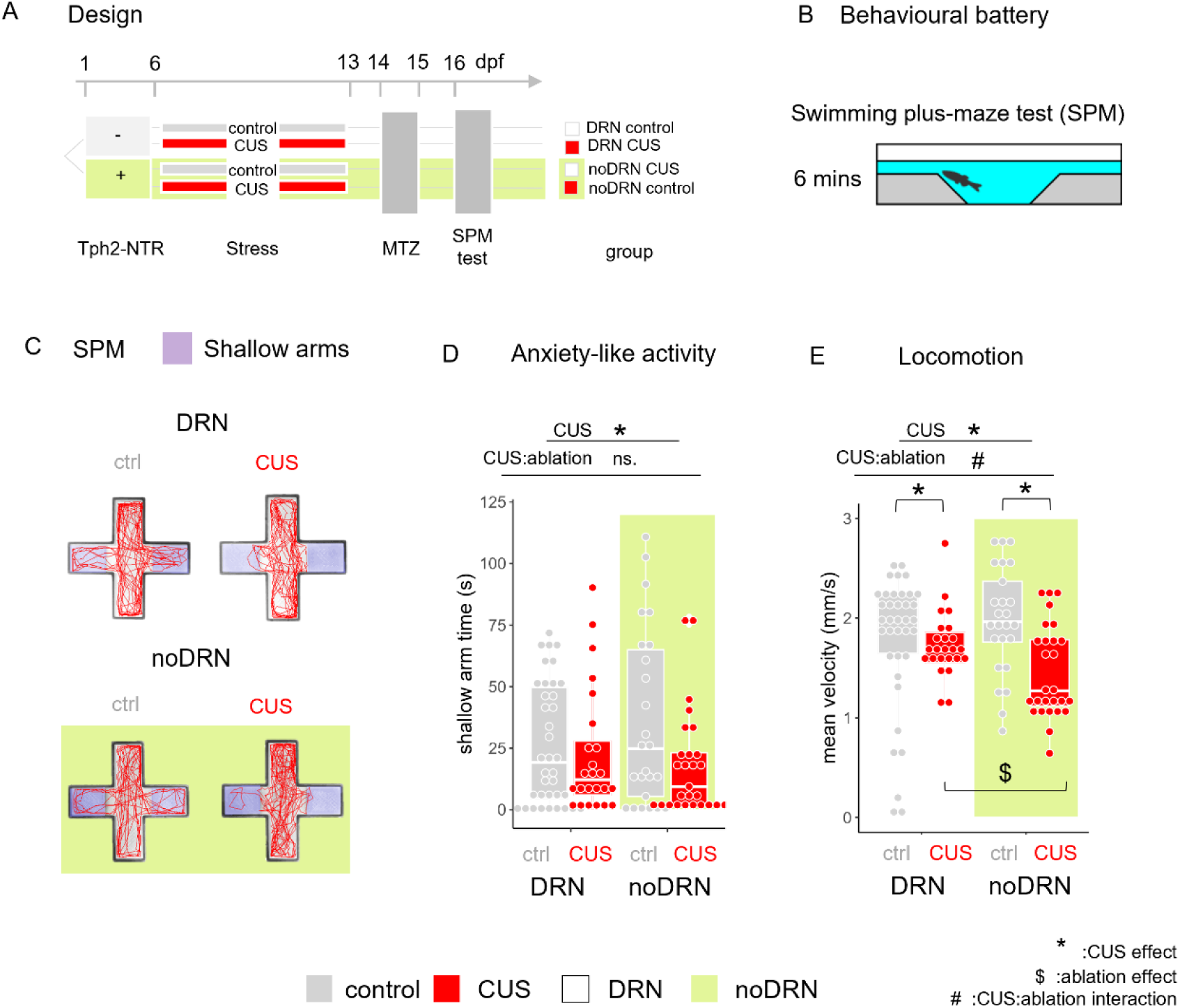
Chronic stress enhances anxiety-like behavior independently from serotonergic DRN function. **A.** Experimental design. Larvae were sorted to nitroreductase (NTR) expressing (+) and non-expressing (−) subjects, then allocated to control or CUS treatment. Metronidazole (MTZ) treatment and the swimming plus-maze (SPM) anxiety assay were done to all subjects **B**. Schematic figures of the SPM test. **C**. Representative swimming trajectories in the SPM from each group. **D**. anxiety-like activity (shallow arm time is negatively correlated with anxiety) and **E**. locomotion (mean velocity) in the SPM. n (DRN ctrl, DRN cus, noDRN ctrl, noDRN cus) = 37, 24, 25, 27. Statistical results for all panels are available in Table 4.

**Table 4:**
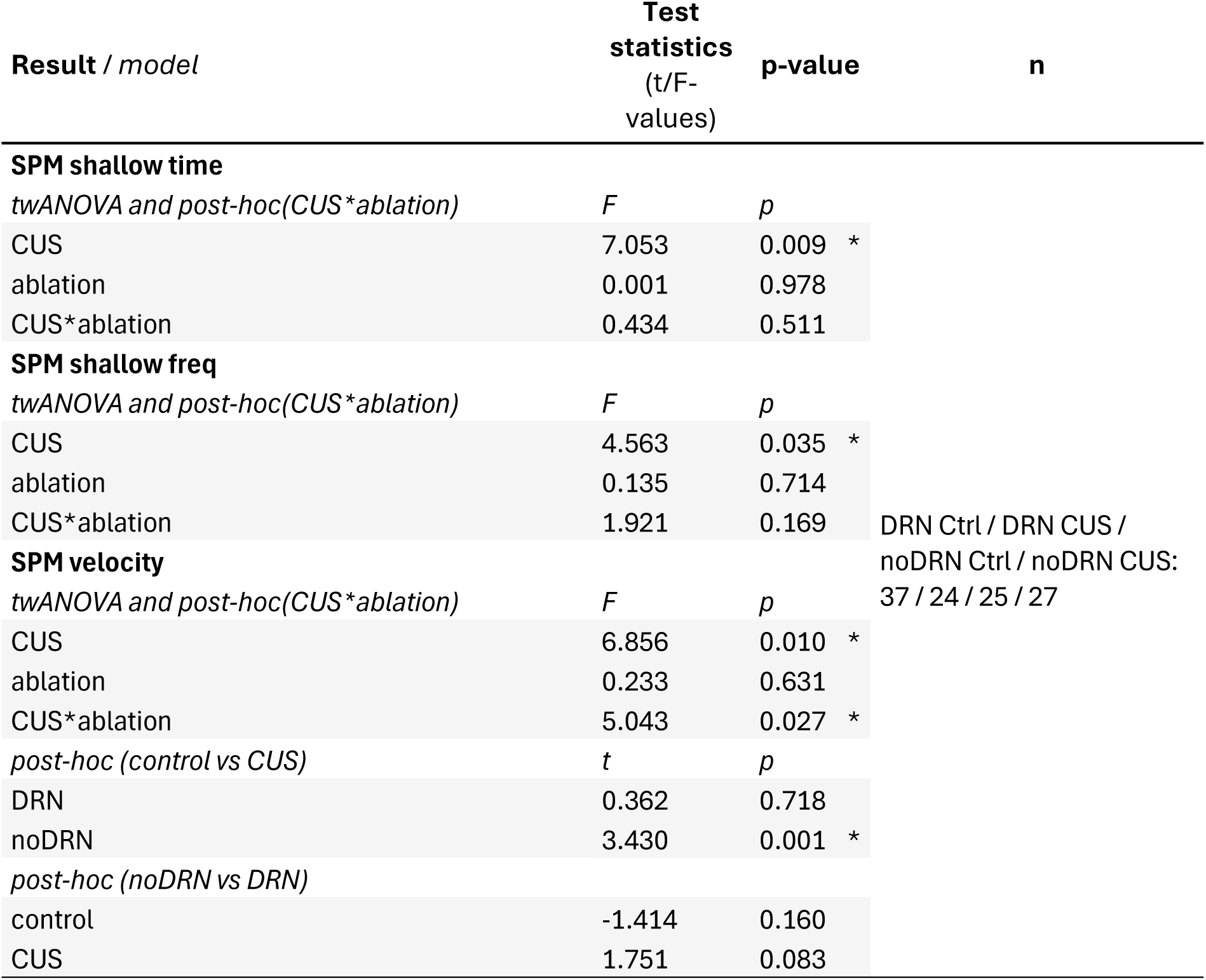
CUS and DRN ablation on anxiety and locomotion.

We found a significant main effect of CUS and no interaction with the ablation on time spent in the shallow arms of the SPM (Fig 4D), which indicated that CUS generally increases anxiety regardless of the ablation.

Interestingly, we found an interactional effect between CUS and ablation on locomotion in the SPM (Fig 4E), similar to that in the habituation assay (Fig 3G). Here, CUS exposure decreased swimming speed in both CUS-exposed groups relative to their respective controls. In addition, DRN ablation further decreased swimming speed when comparing both CUS-exposed groups.

In summary, our results suggest that CUS-induced anxiety is independent from serotonergic DRN function, while the sensitizing effects of the DRN ablation on locomotion are apparent across multiple assay types. Statistical results for all panels are available in Table 4.

## Discussion

### Conserved neuronal composition of the dorsal raphe nucleus after early life stress

There are conflicting findings regarding the effect of prolonged stress exposure on DRN neuroarchitecture in rodents. Some studies found little effect of chronic stress on overall DRN 5-HT neurons abundance ^51,52^, while others reported an increase in the number of DRN Tph2-expressing neurons in rats susceptible to a social challenge after chronic social defeat ^53^. Our results are in line with the former, as the early life chronic stress protocol used here did not alter the general neuronal composition of the developing zebrafish DRN (Supp Fig 2C-D).

Previous studies have observed GABAergic populations of neurons within the zebrafish larvae DRN ^42^. We show here that a significant subset of Tph2 neurons co-express GABAergic markers, representing around 30% of serotonergic DRN neurons in 2 w.o. zebrafish (Supp Fig 2C), a finding paralleled in a recent study of the 1 month-old zebrafish DRN ^54^. Neurons co-expressing serotonin and GABAergic markers are also present in rodents, where they account for 1-17% of the serotonergic neurons depending on the DRN sub-region and the species ^34,55–57^. Interestingly, in mice, exposure to a short-term intense stress induces an increase in the percentage of 5-HT DRN neurons co-expressing GABA ^58^. Here, we did not find evidence that 1-week long CUS exposure changed the ratio of double-positive Tph2-Gad1b neurons in the DRN

### Chronic stress impairs habituation in dorsal raphe serotonergic neurons

Stress exerts complex effects on the serotonergic system, notably because acute short-term stressors or noxious stimuli elicit excitatory responses in 5-HT DRN neurons ^21,42,49,59–61^, whereas chronic or repeated stress decreases 5-HT DRN basal activity in rodents ^23,24,62^. By examining the 5-HT DRN response to an acute stressor in unstressed and chronically stressed fish, our results confirmed and extended these previous studies. We found that most DRN serotonergic neurons were initially strongly excited by hyperosmotic solution, which activates the stress response and elicits robust avoidance in zebrafish larvae ^48,49^. We also found that the responses of DRN 5-HT neurons to repeated hyperosmotic shock rapidly decrease in control fish. Strikingly, this habituation of DRN neurons was disrupted when fish had previously been exposed to chronic stress. Our results are in agreement with prior studies in zebrafish larvae, which found that the activity of the 5-HT DRN decreases within minutes in response to short-term stressor exposure, such as repeated acoustic startles ^22^ or mild electric shocks ^29^. Here, we further demonstrate that chronic stress induces long-term plasticity in the habituation profile of the DRN, which persisted for several days after the last stressor exposure.

We found that impaired DRN habituation following chronic stress was driven by an decreased plasticity of excitatory/inhibitory responses, which was particularly pronounced within serotonergic neurons co-expressing a GABAergic marker. CUS exposure decreased the ratio of plastic- and increased the ratio of stably inhibited response types of Tph2+Gad1b+ cells (Figure 2C-E). This might be due to a different stress susceptibility of subsets of DRN neurons depending on their GABAergic phenotype. In line with this, a study in rodent DRN found that 5-HT/GAD67 neurons possess different electrophysiological properties (lower input resistance and firing rates) than non-GABAergic 5-HT neurons and are more responsive to an anxiogenic environment such as open field exposure ^57^. Interestingly, these stably inhibited Tph2+Gad1b+ neurons show a more caudal distribution in the DRN (Supp Fig 2E), indicating that the CUS facilitated the inhibition of a potentially separate subpopulation of GABAergic cells.

### Chronic stress-induced plasticity in 5-HT DRN neurons drives the disrupted behavioral habituation

At the behavioral level, habituation is a reduction in motor response elicited by repeated presentations of an unexpected stimulus. It is a non-associative form of learning that can be altered in a number of brain disorders in human, including depression and generalized anxiety disorders ^63–65^, and can be modeled with good validity by measuring startle responses in rodents and zebrafish. Here, we found that exposure to 8-day mild unpredictable stress resulted in impaired visual habituation in larval zebrafish. Although it is established that prolonged stress affects other types of learning (e.g. spatial, reversal ^66,67^, the evidence for its effect on habituation learning is scarce. One study found that chronic overexpression of corticotropin-releasing hormone disrupted habituation to acoustic startle in mice ^68^.

A wealth of studies have shown that behavioral habituation is under the control of serotonergic signaling across species ^22,69–71^. In general, high levels of serotonin prevent habituation, whereas lowering serotonin facilitates habituation. In larval zebrafish, Wolman and colleagues found that behavioral habituation to acoustic stimuli is modulated by several serotonergic agents in an unbiased high-throughput drug-screening ^72^. Pantoja and colleagues demonstrated the specific contribution of the DRN to acoustic startle in zebrafish. In their study, decreasing 5-HT content in DRN neurons increased habituation, whereas optogenetic activation of 5-HT DRN reduced habituation ^22^. Varga and colleagues found that in control conditions and proper habituation capacity, the whole-brain 5-HT level is decreasing in response to an exposure to the light/dark test (LDT). In contrast, following chronic social isolation stress, increased central 5-HT is accompanied with decreased habitation in the LDT, an effect that was preventable by the pharmacological decrease of the 5-HT content ^9^. In line with these results, we found that chronically stressed fish displaying altered habituation of 5-HT DRN neurons, which presumably resulted in higher 5-HT levels than in controls, also showed impaired habituation of motor response. Ablating 5-HT DRN neurons did not facilitate habituation compared to control fish with intact DRN. However, it rescued habituation in CUS-exposed fish, thereby causally implicating for the first time the chronic stress-induced plasticity of 5-HT DRN neurons in the decreased habituation capacity.

It is important to note that mammalian 5-HT DRN neurons co-express several neurotransmitters and neuropeptides, including GABA, Glutamate and Dopamine and various neuropeptides ^34,40,73^. Consistent with this, we showed that a third of zebrafish 5-HT DRN neurons co-express GABAergic markers, a finding paralleled in another recent study of the zebrafish DRN ^54^. Interestingly, high throughput drug screening in larval zebrafish recently identified GABAergic signaling as an important modulator of visual habituation learning ^74^. Therefore, it is possible that GABA or other neurotransmitters than serotonin contribute to the role of zebrafish 5-HT DRN neurons in driving stress-induced impairment in habituation.

### Early life chronic stress disrupts behavior both in DRN-dependent and - independent manners

We found that chronic stress impairs behavioral habituation, enhances anxiety-like responses and decreases locomotion. The latter findings are well-established in the literature, as exemplified in two recent meta-analyses emphasizing the maladaptive effects of chronic stress on anxiety and locomotion in rodent ^75^ and in zebrafish models^76^.

Ablating the 5-HT DRN post-exposure to chronic stress in our study further revealed that 5-HT neurons in the DRN are selectively responsible for the disrupted habituation, but not for the increased anxiety. In addition, ablation of the DRN did not change the anxiety levels of unstressed fish in the SPM, reinforcing the notion that 5-HT DRN did not play a role in anxiety-like behaviors measured in our study. The lack of DRN control over CUS-induced anxiety may sound surprising, as the role of DRN in anxiety-like responses is well established in rodent literature. However, extensive reviews of the topic conclude that such correlation depends on the investigated and/or manipulated sub-regions, projections and cell-types of the DRN ^50^. In rodents, separate subsets of DRN neurons are associated with the expression of anxiety-like or depression-like symptoms or with the inhibition of panic-like symptoms ^50,77,78^ and a similar topographical segregation was also found in the zebrafish in response to an anxiogenic environment ^21^ or to aversive stimuli ^54^. In the current study, although our manipulation was restricted to Tph2+ neurons, the ablated cells represent a heterogeneous population, and behavioral differences may arise from this diversity. Consequently, we cannot exclude the possibility that specific subpopulations of DRN serotonergic neurons play distinct roles in anxiety regulation – roles that may not be detectable through our broader manipulation.

The involvement of the DRN in anxiety regulation may also depends on which factors trigger the anxiety-like responses – potentially through the activation of particular neuron subpopulations ^79,80^. The DRN has been implicated in anxiety emerging as a consequence of stress, - in rodents subjected to chronic social defeat ^50,81^ and in zebrafish exposed to repeated mild electric shocks ^29^. In line with this connection between serotonergic function and stress-induced anxiety, social isolation of late larval stage zebrafish was shown to increase serotonergic fiber density, whole-brain serotonin content as well as anxiety-like behavior during novelty challenge ^9^. These results, although support the link between stress, the DRN and anxiety, represent different experimental conditions compared to our study in which anxiety emerged in response to novelty stress following chronic unpredictable stress.

It is also important to note that despite our ablation protocol fully eliminated CUS-impaired 5-HT DRN, it did not eliminate areas that were affected by such impaired DRN activity throughout the CUS period. Consequently, we could only rescue CUS-induced behavioral effects that directly and acutely involved the 5-HT DRN at the time of testing. This does not rule out that 5-HT DRN neurons play a role in the acquisition of CUS-induced maladaptive outcomes during the CUS, but we showed here that the DRN is not involved in the maintenance or the expression of anxiety-like behaviors after the CUS exposure is completed.

We also found that the ablation of the serotonergic DRN reveals, or worsens, CUS-induced hypolocomotion in two different assay settings, i.e. in the SPM and in an open field-like environment during the baseline recording preceding the habituation assay. In both cases, the amplification of CUS-induced hypolocomotion by DRN ablation suggests that adaptation of 5-HT DRN activity during chronic stress might exert a protective role on locomotion. Although the precise neural circuit’s mechanisms underlying this effect are yet to be determined, both anatomical and functional data support that the 5-HT DRN, and serotonin in general, play an important role in adaptively gating locomotion speed depending on recent experience and context ^42,82,83^.

## Acknowledgements

This study was supported by the Research Council of Norway (FRIPRO grant 262698 to FK), and by the Lundbeck Foundation (Ascending Investigator to FK). We thank Astrid Bjørkøy, Head of the Center for Advanced Microscopy (CAM) at the Faculty of Natural Sciences at NTNU for expert guidance with the use of the Leica SP8 SMD/MP.

---

**Supplementary Figure 1:**
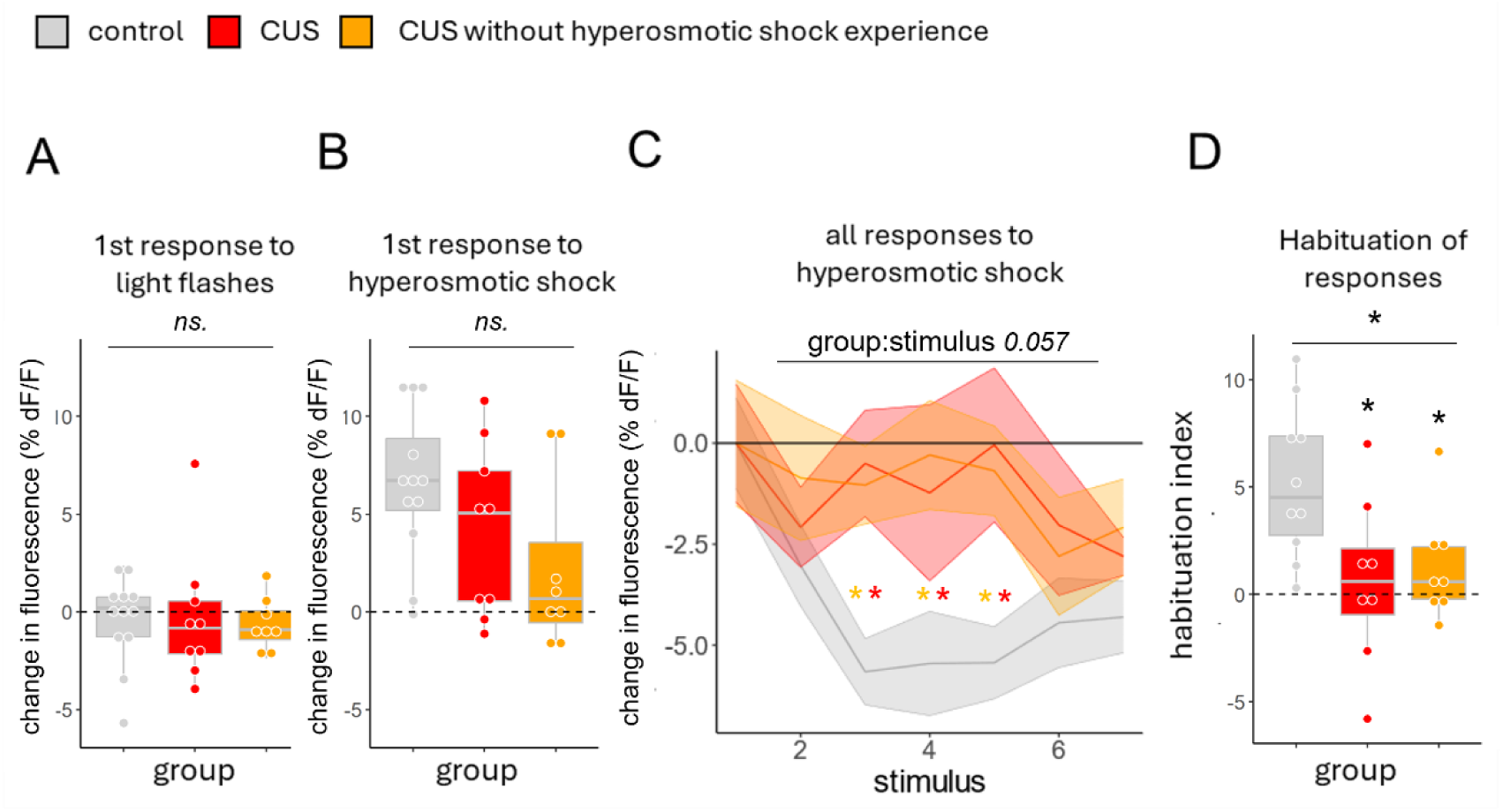
Habituation of DRN responses to hyperosmolarity stimuli in control, and CUS-exposed fish with or without prior hyperosmotic shock experience. **A.** Average DRN neurons response to the 1^st^ light flash. **B.** Average DRN neurons response to the 1^st^ hyperosmolarity stimulus. **C.** Average change in the response amplitude of the DRN to repeated hyperosmotic stimuli relative to the first hyperosmotic stimulus presentation. **D**. Response habituation index. Each dot represents a fish (n (ctrl, cus,cus no osmo) = 12, 9, 8).

**Supplementary Figure 2.**
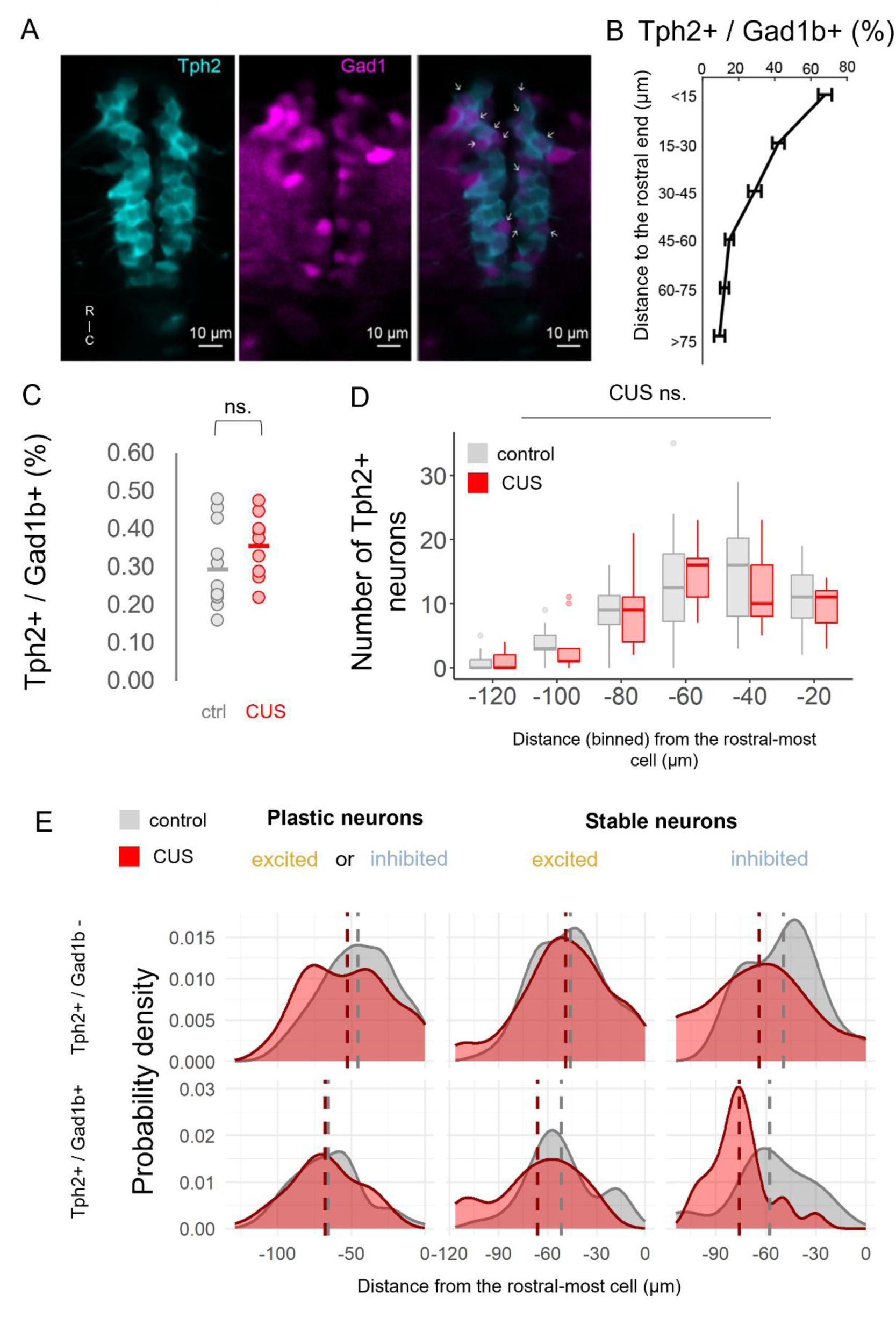
Rostro-caudal distribution of subpopulation of serotonergic neurons within the DRN. **A**. Two-photon image of the dorsal raphe nucleus in a 14 dpf *Tg(tph2:Gal4; uas:GCaMP6s; Gad1b:DsRed)* zebrafish illustrating that a subpopulation of Tph2+ (cyan) neurons, indicated by white arrows, co-expresses Gad1b (magenta). **B.** Percentage of double-positive (Tph2+ / Gad1b +) neurons in the rostro-caudal axis of the DRN calculated in 15 um long regions. **C.** The percentage of double positive neurons among all DRN neurons. **D.** Number of Tph2+Gad1b+ neurons along the rostro-caudal axis of the dorsal raphe nucleus of control and CUS-exposed fish. 0 indicates rostral-most neuron within a fish and negative values go towards caudal. **E.** Density of plastic and stable Tph2+Gad1b+ and Tph2+Gad1b-neurons along the rostro-caudal axis. R:rostral,C:caudal.

**Supplementary Figure 3:**
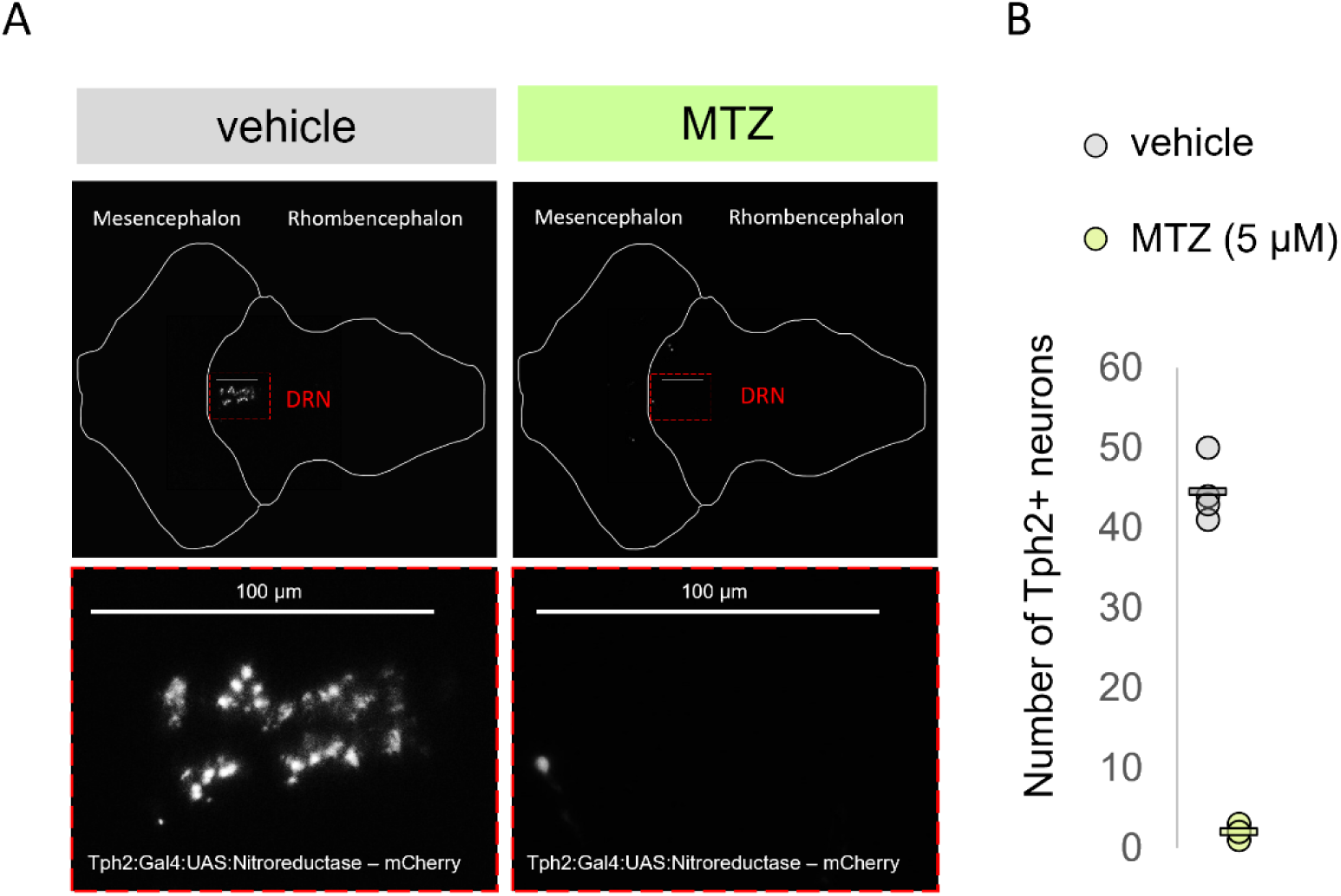
Nitroreductase – metronidazole ablation of serotonergic neurons in the dorsal raphe nucleus. **A.** Representative confocal pictures of intact Tph2+ neurons in the dorsal raphe nucleus of a non-ablated 16 dpf fish (Tph2:Gal4:UAS:Ntr-mCherry+ exposed to vehicle, left) and near complete lack of Tph2+ neurons in an ablated fish (Tph2:Gal4:UAS:Ntr-mCherry+exposed to metronidazole, right). **B.** Number of Tph2+ neurons in 4 vehicle vs 4 MTZ-treated fish.

**Supplementary Figure 4:**
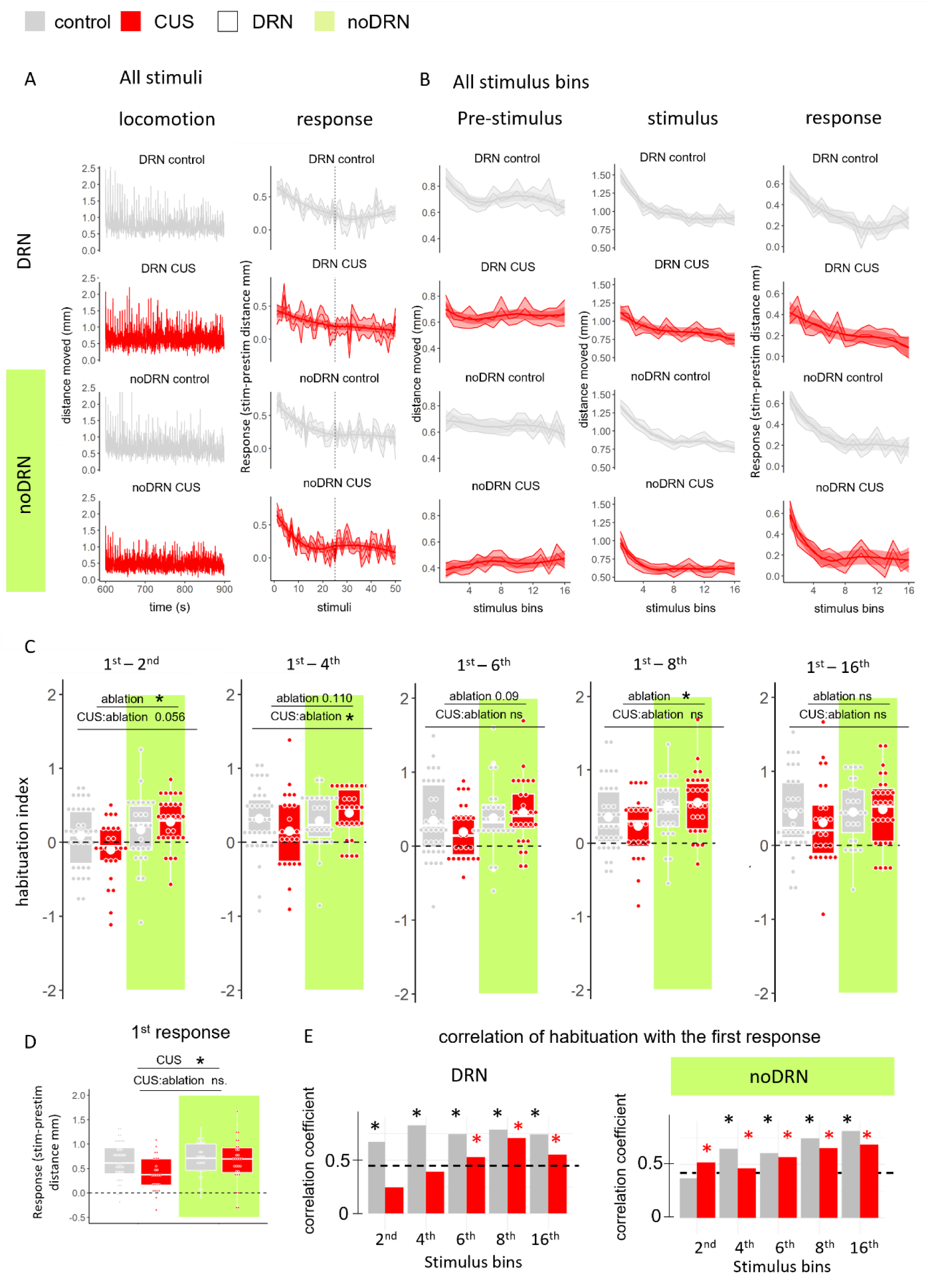
Overview of basal and stimulus evoked locomotion in the dark flash habituation assay. **A.** Distance moved (left) and dark flash response right) during the whole stimulus period. Vertical dashed line on the right panel indicates the 25^th^ stimulus, i.e. the middle of the test. Note that habituation already happened to this point in all but not in the CUS-exposed, intact DRN group. **B**. Distance moved pre-stimulus(left), during stimulus (middle) and stimulus responses (during – pre stimulus distance)(right) in all response bins. Habituation trends remain intact, but variability decreased in binned data. **C**. Habituation indices at the 2^nd^, 4^th^, 6^th^, 8^th^ (half of test) and the 16^th^ (end of test) stimulus bins. Horizontal dashed lines were added to indicate no habituation. **D**. Response to the first stimulus. Horizontal dashed line at 0 indicates no response. **E**. Pearson correlation coefficients between the first and the 2^nd^, 4^th^, 6^th^, 8^th^ and 16^th^ stimulus bins. Black or red asterisk indicates significant correlation following control or CUS treatment, respectively. Correlation coefficients from this analysis were all significant above the horizontal dashed lines. n (DRN ctrl, DRN cus, noDRN ctrl, noDRN cus) = 38, 27, 27, 29.

**Supplementary Table 1:**
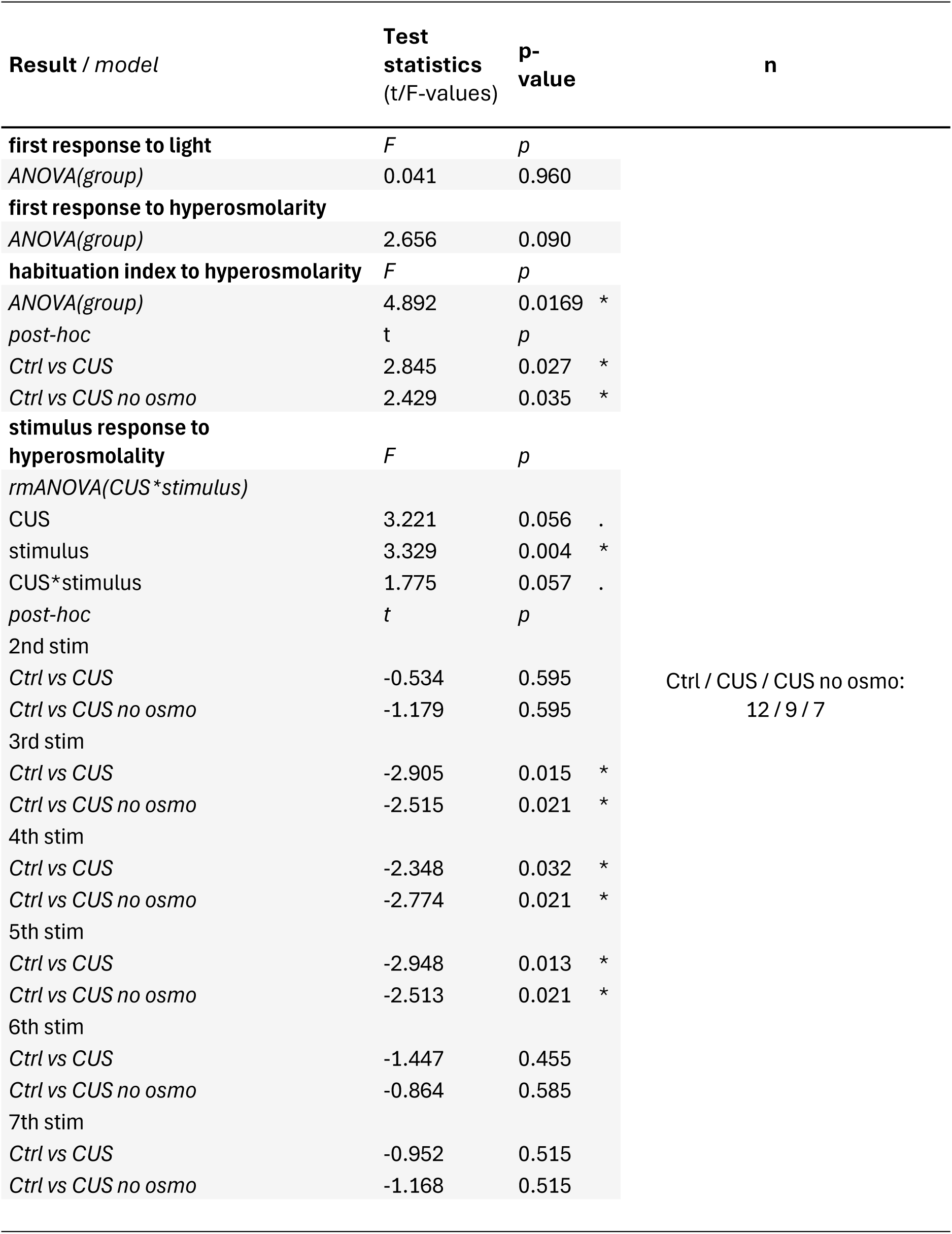
CUS on DRN activation.

